# Reactive Oxygen Species Mediate Transcriptional Responses to Dopamine and Cocaine in Human Cerebral Organoids

**DOI:** 10.1101/2023.06.13.544782

**Authors:** Thomas T. Rudibaugh, Albert J. Keung

## Abstract

Dopamine signaling in the adult ventral forebrain regulates behavior, stress response, and memory formation and in neurodevelopment regulates neural differentiation and cell migration. Excessive dopamine levels including due to cocaine use both in utero and in adults could lead to long-term adverse consequences. The mechanisms underlying both homeostatic and pathological changes remain unclear, partly due to the diverse cellular responses elicited by dopamine and the reliance on animal models that exhibit species-specific differences in dopamine signaling. To address these limitations, 3-D cerebral organoids have emerged as human-derived models, recapitulating salient features of human cell signaling and neurodevelopment. Organoids have demonstrated responsiveness to external stimuli, including substances of abuse, making them valuable investigative models. In this study we utilize the Xiang-Tanaka ventral forebrain organoid model and characterize their response to acute and chronic dopamine or cocaine exposure. The findings revealed a robust immune response, novel response pathways, and a potential critical role for reactive oxygen species (ROS) in the developing ventral forebrain. These results highlight the potential of cerebral organoids as *in vitro* human models for studying complex biological processes in the brain.

## Introduction

Dopamine signaling in the ventral forebrain is crucial in regulating behavior, stress response, and memory formation in adults^1–4^. Additionally, it holds a significant role in neural differentiation and cell migration during neurodevelopment^5, 6^. Excess levels of extracellular dopamine can lead to adverse long-term consequences, such as addiction formation, memory loss, premature neural differentiation, and neuroinflammation^5, 7–14^. Despite its importance, the mechanisms underlying these alterations remain unclear, and in particular, our understanding of dopamine signaling in human systems is even further limited^12, 14–1617–19^.

Previous studies have shown human pluripotent stem cells can be differentiated in 2-D towards ventral forebrain neurons expressing functional dopamine receptors^20–25^, and these neurons can even recapitulate complex phenomena such as a gene desensitization following chronic dopamine exposure^26^. Recently, 3-D cerebral organoids have been developed to model neurodevelopment, cell migration patterns of different brain regions^27, 28^, and stem cell and neuronal responses to external stimuli, including substances of abuse^29–32^. It will be important to characterize organoid responses more deeply to stimuli, to help identify underlying pathways involved in dopamine function, to determine the phenotypes that organoids can replicate well, and identify those areas for further improvement.

In this work, we leverage the previously described Xiang-Tanaka ventral forebrain organoids to model ventral forebrain neurodevelopment and investigate their responses to acute and chronic dopamine exposure using a combination of next generation sequencing and electrophysiological measurements^28^. Through our analysis, we observe immune related transcriptional responses and identify novel response pathways activated following dopamine exposure. Using small molecule inhibitors, responses are found to act at least in part through reactive oxygen species and in part through dopamine receptor signaling. Finally, we observe some similarities in the responses to chronic cocaine exposure. These findings demonstrate the utility of cerebral organoids as *in vitro* human platform for studying complex biological processes and their potential mechanisms.

## Results

### Ventral forebrain organoids exhibit Ca^2+^ and intracellular cAMP responses to dopamine

Xiang-Tanaka and colleagues described the generation of human medial ganglionic eminence organoids (hMGEOs) through the activation of the sonic hedgehog (SHH) pathway to achieve ventral forebrain identity^28^. In this work, we characterize the functional and transcriptomic responses of these organoids to dopamine and cocaine. We first confirm the generation of ventral forebrain identity. By day 30 (D30), *GSX2*+/*SOX2*+ progenitor cells arise, suggesting an early ventral forebrain fate^21, 33^ (Figure 1A). By D90, we observe robust expression of *GABA*+/*MAP2*+/*CTIP2*+, indicating development of GABAergic neurons (Figure 1B), and of medium spiny neuron (MSN) marker *DARPP32* (Figure 1C). After confirming a ventral forebrain fate, we assess the presence of electrophysiological activity, observing spontaneous action potentials through Fluo-4 Ca^2+^ imaging in D90 organoids (Figure S1A). These transients are inhibited by the addition of the sodium channel blocker tetrodotoxin (TTX), indicating the Ca^2+^transients are likely dependent on neuronal activity. Taken together, these results show the differentiation of stem cells into relatively mature and electrophysiologically active ventral forebrain organoids^28^.

**Figure 1.**
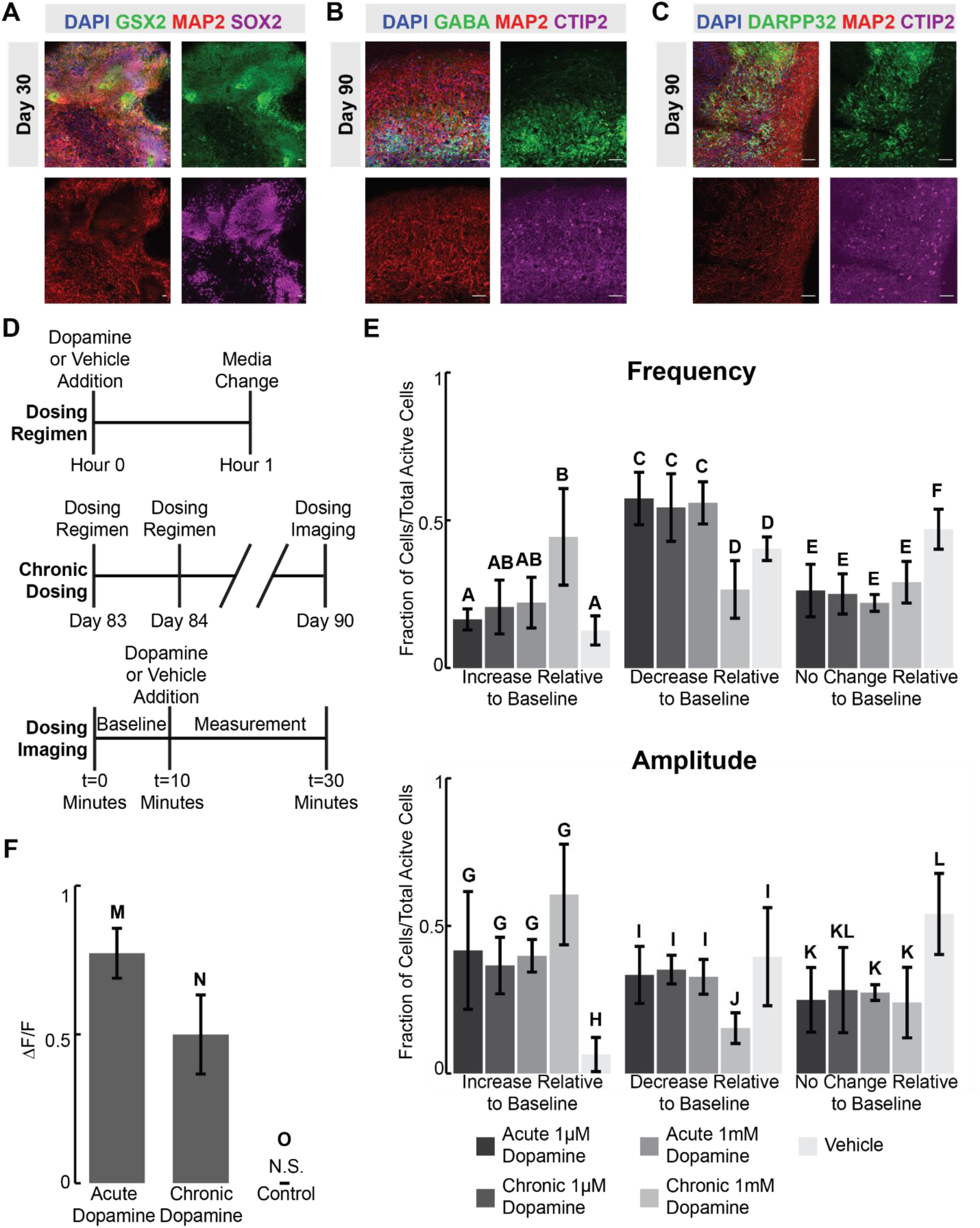
Ventral forebrain organoids exhibit Ca^2+^ and intracellular cAMP responses to dopamine A-C) Immunostaining images of **A)** day 30 or **B-C)** day 90 (D90) Xiang-Tanaka organoids demonstrating expression of relevant ventral forebrain markers. **D)** Schematic illustrating the dosing regimen and chronic dosing conditions used for Ca^2+^ and cAMP imaging experiments. **E)** Ca^2+^ imaging frequency and amplitude measurements of individual neurons compared to their baseline controls following acute or chronic 1µM or 1mM dopamine exposure. An increase or decrease was defined as a cell having greater than 20% deviation from its baseline measurement following compound exposure. **F)** Amplitude of intracellular cAMP change relative to baseline following acute or chronic 1µM dopamine exposure. **E-F)** Error bars represent 95% confidence intervals for n=3 technical replicates. * p<0.05, one-way ANOVA with Tukey Kramer post hoc analysis.

Dopamine is a major neurotransmitter impinging on ventral forebrain neurons. In previous work, 2-D ventral forebrain neurons responded electrophysiologically and transcriptionally to dopamine^5, 6, 24, 26^. In addition, there is evidence to suggest cultured neurons can exhibit neuroplastic alterations following chronic dopamine exposure^26^. We ask if this organoid model similarly exhibits a response to dopamine and if this response changes following chronic dopamine exposure. It is unclear, however, what dopamine concentration correlates to *in vivo* concentrations at the synaptic cleft. Previous work in mice suggests dopamine concentrations anywhere from 100nM-100μM, while human cell culture methods have used higher concentrations of up to 10mM dopamine to elicit a response^26, 34–36^. Therefore, we treat D90 organoids to either an acute or chronic dose of 1μM or 1mM dopamine along with a vehicle control (Figure 1D-E, Figure S1B). Surprisingly, neither dopamine concentration nor chronic exposure causes a change in firing frequency or amplitude when measured at the population level (Figure S1B).

One possible explanation is that dopamine is causing opposing responses in different cell types that average out at the population level. D1 type dopamine receptor stimulation usually causes an increase in action potential frequency whereas D2 receptor stimulation usually leads to a decrease, although there are exceptions^37–39^. While it is technically challenging to track Ca^2+^ transients simultaneously with identifying the neuronal subtype of each cell, we can longitudinally compare Ca^2+^ transient frequency and amplitude of each individual cell before and after dopamine or vehicle control exposure. We limit our analysis to the active cells who exhibit at least one measurable calcium transient during the 10-minute baseline measurement. The results show over 50% of cells treated with vehicle controls have the same firing frequency and amplitude as their baseline, while only 20-30% of dopamine treated cells maintain the same firing frequency and amplitude (Figure 1E). This suggests that of the cells measured, roughly 25% are exhibiting a change in Ca^2+^ transients that can be directly attributed to dopamine. Both acute 1µM and 1mM dosing regimens lead to a similar phenotype, where many individual cells exhibit a decrease in Ca^2+^ transient frequency and an increase in amplitude following dopamine exposure. In addition, we observe a robust response following 1μM dopamine exposure, suggesting this lower concentration is enough to drive a cellular response in organoids.

In addition to tracking Ca^2+^ transients, we can also assess how intracellular cAMP changes in response to dopamine. Dopamine signaling is known to drive either an increase or decrease in intracellular cAMP depending on the neural subtype^12, 40–42^, and cAMP leads to a transcriptional response in ventral GABAergic neurons, ultimately contributing to many of the downstream neuroplastic changes associated with chronic drug exposure^3, 10, 43, 44^. To our knowledge, cAMP and complex downstream responses have not been previously described in stem cell-derived systems. Thus, we expose organoids to acute and chronic 1µM dopamine and measure the levels of intracellular cAMP (Figure 1D & 1F, Figure S1C). Reassuringly, dopamine causes an increase in the amount of intracellular cAMP, and this response is dampened in the chronically dosed samples (Figure 1F, Figure S1C), indicating a potential desensitization phenotype.

### Transcriptional analysis reveals immune-related response to acute and chronic dopamine

While Ca^2+^ and cAMP measurements indicate functional responses to dopamine, transcriptomic changes could provide deeper insights into how longer term neuroplastic alterations might arise^8, 14, 45, 46^. In addition, transcriptomic changes caused by chronic dopamine overexposure are implicated in addiction formation in adults and neurodevelopmental alterations *in utero*^5, 8, 47, 48^. Therefore, we expose D90 organoids to acute and chronic doses of 1μM dopamine followed by bulk RNA sequencing and differential gene expression (DGE) analysis relative to D90 vehicle control (Figure 2, Figure S2, Supplemental Excel File 1). Chronic dopamine exposure causes an increase in the number of differentially expressed genes (DEGs) compared to the acutely dosed samples (Figure 2B). Among the top 20 downregulated genes in the acutely dosed samples are many genes in the HOX protein family, which play a role in cell migration and apoptosis during neurodevelopment^49, 50^ (Figure 2C, left). Among the top upregulated DEGs in the acutely dosed organoids are the anti-inflammatory cytokines, *IL1R1* and *TGFβI,* along with two genes linked with a pro-inflammatory immune response, *FCN1* and *SPON2.* In addition, we observe activation of the ECM-related genes *SERPINE1*, and *FN1*^51–57^. Among the top 20 upregulated genes for the chronically dosed organoids are pro-inflammatory immune response markers, *S100A9* and *FGR*, along with anti-inflammatory immune response markers, *HSD11B1* and *ITGB2*^58^ (Figure 2C, right). In addition, we observe upregulation of multiple collagen proteins.

**Figure 2.**
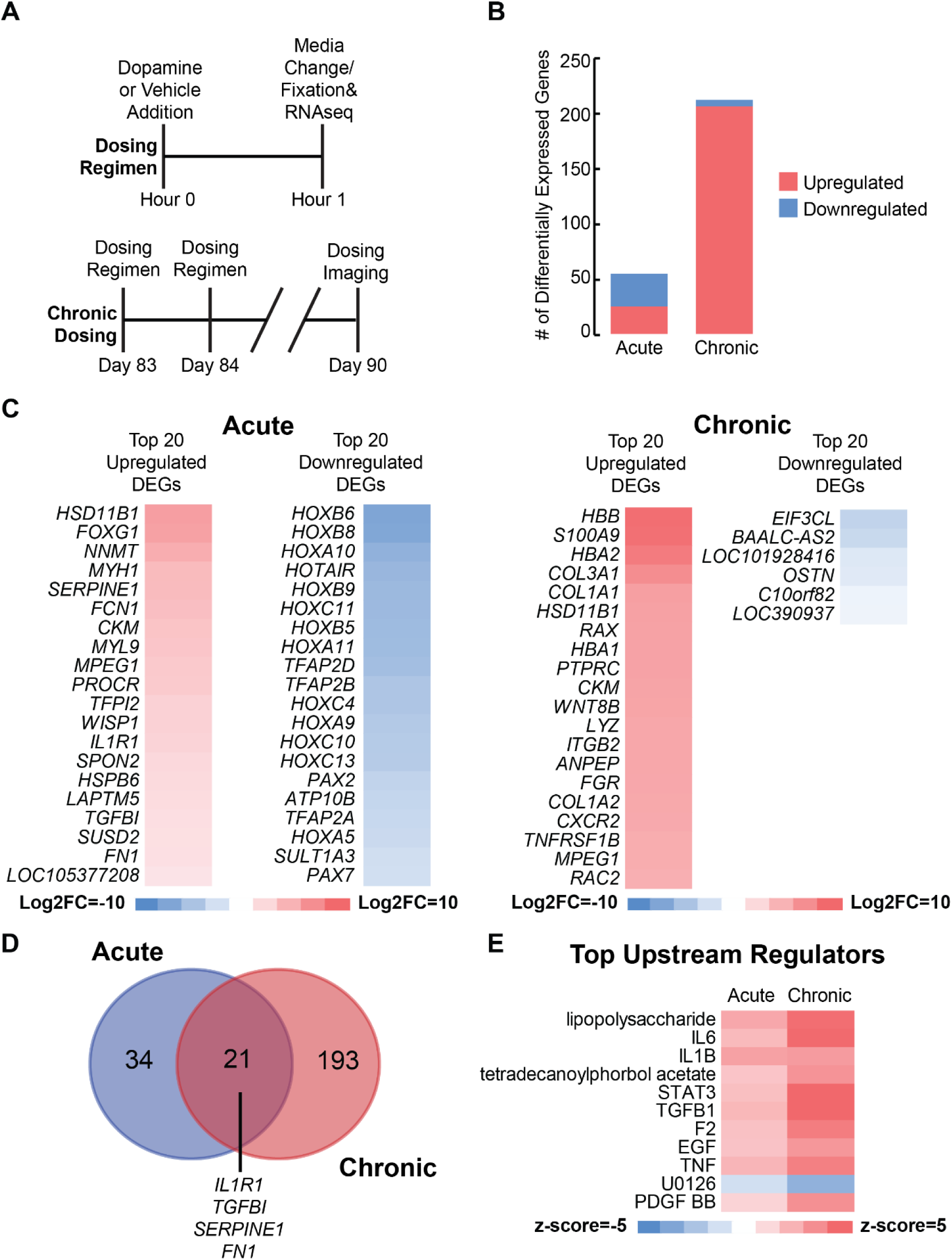
Transcriptional analysis reveals immune-related response to acute and chronic dopamine. **A)** Schematic illustrating the dosing regimen and chronic dosing conditions used for bulk sequencing experiment. **B)** Counts of differentially expressed genes (DEGs) following acute and chronic dopamine exposure. Differential gene expression (DGE) performed relative to vehicle controls. Genes are considered differentially expressed when they have log2FC>|1| and false discovery rate q<0.05. **C)** Heat map of the top 20 up and down regulated genes following acute or chronic dopamine exposure. **D)** Venn diagram showing the overlap between the acute and chronic DEGs. **E)** Heat map of the top ten regulated upstream regulators based on z-score using gene set enrichment analysis (GSEA). Regulator up or downregulation is considered significant if z-score>|2|.

Comparison of the DEGs from the acute and chronic conditions shows 21 genes shared between both conditions, and many of these genes are again linked with the ECM and immune response including *IL1R1, TGFβI*, *SERPINE1*, and *FN1* among others (Figure 2D, Supplemental Excel File 2). Gene set enrichment analysis (GSEA) also suggests both the acute and chronic exposures are acting through similar upstream regulators related to either a pro-inflammatory immune response including lipopolysaccharide, *IL6, IL1B,* and *TNF* or through an anti-inflammatory immune response including *TGF-β1, EGF,* and *STAT3*^59, 60^ (Figure 2E).

### Organoid response to dopamine driven by reactive oxygen species

Alterations in the expression of immune-related genes as well as HOX genes suggested dopamine may not only be acting through canonical neuronal signaling pathways. In particular, dopamine has been shown to activate cellular immune responses by both binding to dopamine receptors directly and through dopamine metabolism by monoamine oxidases^12, 13^. As another example, dopamine metabolism by microglia leads to an increase in both intra- and extra-cellular reactive oxygen species (ROS), which in turn leads to oxidative stress and activation of immune cells^61, 62^. This immune response induces ECM remodeling, including the increased expression of multiple collagen proteins especially collagen I proteins, similar to what is observed in our samples^63^. Therefore, we hypothesize that ROS may be mediating the transcriptional responses we are observing to dopamine.

To test this hypothesis, we measure the endogenous ROS levels of D90 organoids exposed to acute and chronic dopamine. Both samples show higher levels of endogenous ROS (Figure 3A). We also measure the levels of cAMP following dopamine exposure in the presence of either the dopamine receptor antagonists SCH-23390 and sulpiride or the ROS inhibitor acetylcysteine (Figure 3B). We observe dopamine receptor antagonists show some ability to blunt the increase in intracellular cAMP while the ROS inhibitor dramatically reduces the intracellular cAMP response. Similarly, we observe that acetylcysteine also substantially blunted the transcriptomic response of 1μM dopamine down to control levels in D90 organoids (Figure 3C-F, Figure S3, Supplemental Excel File 3). To ensure acetylcysteine alone is not driving this phenotype, we included controls both with and without acetylcysteine exposure (Figure 3D).

**Figure 3.**
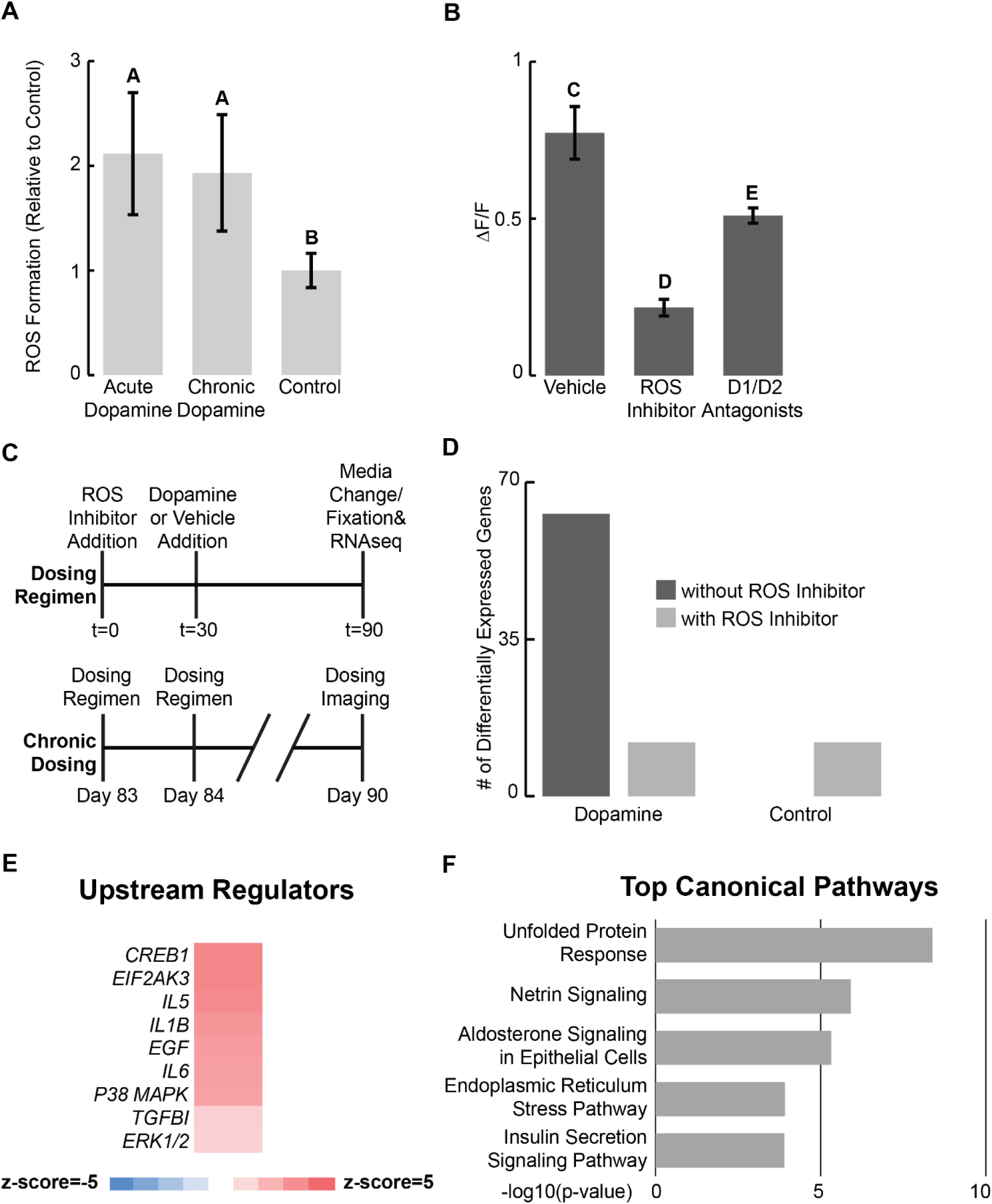
Organoid response to dopamine driven by reactive oxygen species. **A)** Reactive oxygen species in ventral forebrain organoids following acute and chronic dopamine exposure. Values normalized to vehicle controls. **B)** cAMP measurement in ventral forebrain organoids following treatment with either vehicle, an ROS inhibitor, or D1/D2 dopamine receptor antagonists prior to acute dopamine exposure. **A-B)** Error bars represent 95% confidence for n=3 biological replicates. * p<0.05, one-way ANOVA with Tukey Kramer post hoc analysis. **C)** Schematic illustrating the dosing regimen and chronic dosing conditions used for bulk sequencing experiment. Time (t) in minutes. **D)** Number of DEGs following chronic dopamine exposure with or without an ROS inhibitor added 30 minutes prior to dopamine exposure. Differential gene expression performed relative to vehicle controls. Control with an ROS inhibitor added was compared against control without an ROS inhibitor. The dopamine dosed sample without an ROS inhibitor added was compared against the control without an ROS inhibitor added. The dopamine dosed sample with an ROS inhibitor added was compared against the control sample with an ROS inhibitor added. **E)** Heat map showing activation of select upstream regulator immune response genes calculated by z-score using GSEA. Regulator up or downregulation is considered significant if z-score>|2|. **F)** Top 5 regulated canonical pathways by -log10(p-value) identified using GSEA.

GSEA of the DEGs from the chronically dosed samples once again reveals that, among the top upstream regulators are multiple pro- and anti-inflammatory cytokines including *IL5, IL6, EIFAK3*, and *TGFβI* among others^64, 65^ (Figure 3E). In addition, among the top 5 regulated pathways are the unfolded protein response pathway and endoplasmic reticulum stress response pathways (Figure 3F). One key signature of oxidative stress is an increase in unfolded proteins accumulating in the ER^66, 67^. Collectively, these results suggest a potential influence of dopamine on neurodevelopment through ROS.

### ROS also mediates the transcriptomic response to cocaine

Cocaine is related to dopamine signaling in that it causes an increase of dopamine in the synaptic cleft through inhibiting the presynaptic dopamine transporter^8, 47^. Interestingly, prenatal cocaine exposure has also been linked to oxidative stress in the cerebral cortex, causing premature neural differentiation^29, 32, 66, 68^. There is evidence to suggest cocaine acts through a similar mechanism in the developing ventral forebrain as well ^69^. Therefore, we expose D90 ventral forebrain organoids to chronic doses of 1μM cocaine with and without acetylcysteine (Figure 4, Figure S4, Supplemental Excel 4). The presence of acetylcysteine dramatically reduces the number of DEGs (Figure 4B). Among the top upregulated genes following cocaine exposure, we observe upregulation of multiple HOX family genes (Figure 4C). In addition, we also observe upregulation of multiple immune-related genes including *IL1R1*, *SERPINA3*, and *CYP1B1* (Figure 4D)^70^. GSEA suggests many of the upstream regulators identified following dopamine exposure are also activated by cocaine, and this activation is ROS dependent (Figure 4E).

**Figure 4.**
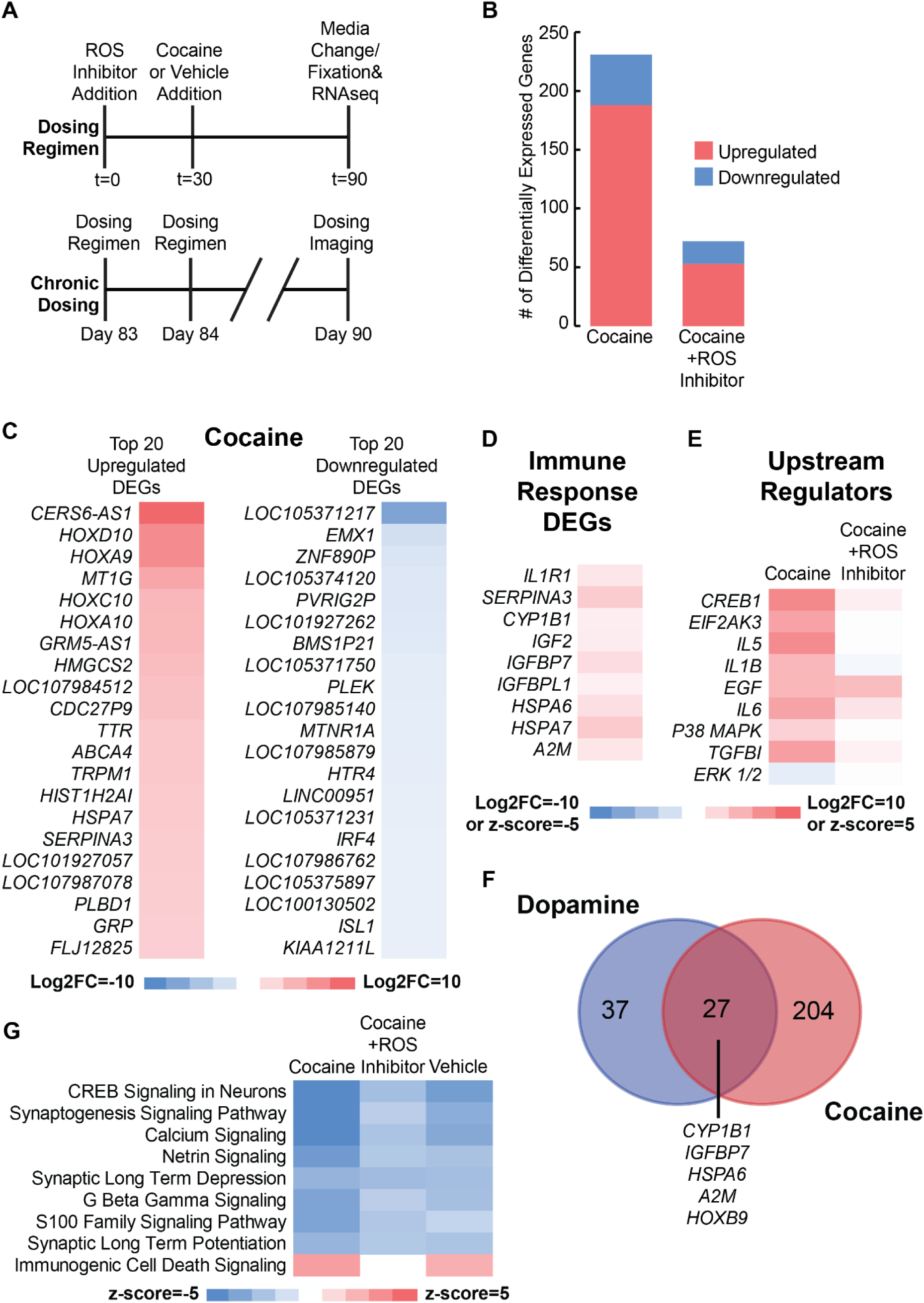
ROS also mediates the transcriptomic response to cocaine. **A)** Schematic illustrating the dosing regimen and chronic dosing condition used for bulk sequencing experiment. **B)** Count of DEGs following chronic cocaine exposure with or without an ROS inhibitor. Differential gene expression performed relative to vehicle controls. The cocaine dosed sample without an ROS inhibitor added was compared against the control without an ROS inhibitor added. The cocaine dosed sample with an ROS inhibitor added was compared against the control sample with an ROS inhibitor added. with or without an ROS inhibitor. **C)** Heat map of top 20 up and downregulated DEGs following cocaine exposure without an ROS inhibitor. **D)** Heat map of immune response related genes identified in figure 3 following cocaine exposure without an ROS inhibitor. **E)** Heat map of upstream regulators identified in figure 3 following cocaine exposure with or without an ROS inhibitor. **F)** Venn diagram showing the overlap between the DEGs following dopamine or cocaine exposure without an ROS inhibitor. **G)** Heat map of the top regulated canonical pathways by z-score following dopamine or cocaine exposure with or without an ROS inhibitor.

Given similarities in ROS mediating transcriptomic responses to dopamine and cocaine, we ask how similar the organoids respond transcriptomically to these two compounds. A comparison of DEGs reveals almost 50% of the DEGs in the dopamine dosed samples are present in the cocaine dosed samples, and many of them are immune-related genes (Figure 4F, Supplemental Excel File 2). GSEA indicates many of the top regulated pathways are the same for both the cocaine and dopamine dosed samples, and ROS inhibition dampens these pathways (Figure 4G). Furthermore, organoids demonstrate downregulation of netrin signaling, responsible for ECM organization and cell migration during neurodevelopment, in both our dopamine and cocaine dosed samples. We also observe upregulation of the immunogenic cell signaling pathway in the cocaine and dopamine dosed samples, which is associated with stress induced apoptosis (Figure 4G). These pathways are both reduced in the acetylcysteine conditions, further suggesting the immune-related and ECM responses caused by dopamine and cocaine are ROS mediated. A similar trend is observed in the upstream regulators (Figure S4B).

## Discussion

Cerebral organoids provide a valuable human model for studying responses to external stimuli^19, 71, 72^. In this study, we demonstrate that ventral forebrain organoids exhibit Ca^2+^, cAMP, and transcriptional responses to dopamine and cocaine^28^. Our results reveal that dopamine and cocaine induce these responses through ROS-mediated mechanisms, and that these responses are blunted upon ROS inhibition. In addition, transcriptomic analyses implicate immune, inflammation, ECM, and HOX genes in these responses.

While it has been widely accepted that dopamine’s effects are primarily driven by dopamine receptor binding and subsequent intracellular signaling in adult neurons, recent evidence from mouse studies suggests that dopamine-mediated reactive oxygen species (ROS) also contribute significantly to neurodevelopment^61, 73, 74^, and that dopamine may regulate cell cycles and differentiation of stem and progenitor cells^5^. Our results using human cerebral organoids suggest that dopamine mediated ROS may hold a similar role in human neurodevelopment, although these findings would need to be validated *in vivo*.

Human stem cell derived models are known to be heterogeneous, and therefore a limitation of this study is the lack of cell type specific resolution. However, this study provides results that suggest several interesting avenues for investigation. First, the transcriptomic responses involving immune, inflammation, ECM, and HOX genes suggest that dopamine and cocaine likely act on multiple different cell types ranging from neurons and microglia to neural stem and progenitor cells, and therefore future work should begin to dissect these different effects in both cell culture and animal models. Second, our results indicate this complexity extends to multiple mechanisms of action, where dopamine can act through dopamine receptors but also through other as yet unknown pathways that may include metabolism or oxidative stress induced by byproducts of dopamine and cocaine.

Third, there are specific pathways and phenotypes that might be explored based upon our results. For example, among the top regulated pathways responsive to dopamine was the downregulation of multiple synapse related pathways including synaptogenesis signaling and synaptic long-term potentiation. Additionally, we observe downregulation of CREB signaling and calcium signaling. Notably, previous research has associated prenatal cocaine exposure with reduced synapse formation^66, 75, 76^. As another example, premature cocaine exposure has been linked with decreased cell migration *in utero* and this response has been associated with downregulation of *BDNF*^54, 75, 77^. Interestingly, our GSEA identified *BDNF* as downregulated in cocaine dosed organoids (Figure S4C)^75, 78^. As a final example, there is an intriguing link between the ECM and immune/inflammation related genes that we observe altered by dopamine exposure. One possible explanation for increased collagen expression is activation of anti-inflammatory macrophages, which when activated leads to ECM deposition^798081^. Given the ability of organoids to generate multiple cell types, this model offers an opportunity to understand these potential cell type specific responses as well as their collective interactions.

### Experimental Procedures hESC Cell Lines

H9 and H1 hESCs (WA09 and WA01; WiCell) were grown in E8 media (Thermo Fisher Scientific) in 6 well culture dishes (Greiner Bio-One) coated with 0.5µg/mL Vitronectin (Thermo Fisher Scientific). Cells were passaged every 3-5 days as necessary using 0.5mM EDTA (Thermo Fisher). All staining and qPCR experiments included in Figure 1 were conducted in H1 and H9 stem cells. Due to cost limitations, all sequencing experiments were limited to H9 stem cells.

### Organoid Culture

The Xiang ventral forebrain protocol organoids were generated as previously described with small modifications^28^. Stem cells were grown to 75% confluency before dissociation into a single cell suspension using EDTA and Accutase (Fisher). 9,000 cells were plated into a low attachment U-bottomed 96 well plate(VWR) in Induction media supplemented with 50µM Y-27632(LC Labs) and 5% v/v heat-inactivated fetal bovine serum (Corning). Induction media contained DMEM-F12(Gibco), 15%v/v knockout serum replacement (Thermo Fisher), 1% v/v MEM-NEAA(VWR), 1% v/v Glutamax(Gibco), 7µL/L β- mercaptoethanol(Amresco), 100nM LDN193189 (Sigma), 10μM SB431452 (Abcam), and 2μM XAV939 (Sigma). After 48 hours, half of the media was replaced with Induction media containing Y-27632 and without heat-inactivated FBS. After another 48 hours, half the media was again removed and replaced with Induction media without Y-27632. From days 6-10 organoids were cultured in a 37°C tissue culture incubator with 5% CO2 and received regular half media changes every 48 hours.

After day 10, organoids were transferred to a deep bottom tissue culture treated 10cm plate (Thermo Fisher) containing differentiation media supplemented with 100ng/mL SHH (VWR) and 1µM purmorphamine (VWR). Differentiation media contained DMEM-F12, 0.15% w/v Dextrose (Sigma), 7µL/L β-mercaptoethanol, 1% v/v N2 Supplement (Thermo Fisher), 1% v/v Glutamax, 0.5% v/v MEM-NEAA, 0.025% insulin solution (VWR), and 2% v/v B27 supplement without vitamin A (Thermo Fisher). Cell culture plates were placed on a shaker in the tissue culture incubator and rotated at 70rpm/min. On day 14 media was removed and replaced with fresh differentiation supplemented with SHH and purmorphamine. On day 18 differentiation media was removed and replaced with maturation media containing vitamin A. Maturation media consisted of a 1:1 mixture of DMEM-F12 and Neurobasal media (VWR), along with 0.5% v/v N2 supplement, 1% v/v B27 supplement with vitamin A (Thermo Fisher), 1% v/v Glutamax, 0.5% v/v MEM-NEAA, 0.025% v/v human insulin solution, 3.5µL/L β-mercaptoethanol, and 1% v/v Penicillin/Streptomycin (VWR). In addition, 20ng/mL of BDNF (Peprotech) and 20ng/mL GDNF (Peprotech) were added to the media. From day 18, organoids were cultured on the orbital shaker with weekly media changes in maturation media.

### Cryosectioning and Immunohistochemistry

Tissues were fixed in 4% paraformaldehyde (Sigma) for 15 minutes at 4C followed by 3, 10-minute PBS washes (Gibco). Tissues were placed in 30% sucrose overnight at 4C and then embedded in 10% gelatin/7.5% sucrose (Sigma). Embedded tissues were flash frozen in an isopentane (Sigma) bath between −50 and −30C and stored at -80C. Frozen blocks were sectioned (Thermo Fisher) to 30-µm. For immunohistochemistry, sections were blocked and permeabilized in 0.3% Triton X-100 (VWR) and 5% normal donkey serum (VWR) in PBS. Sections were incubated with primary antibodies in 0.3% Triton X- 100, 5 % normal donkey serum in PBS overnight at 4C. Sections were then incubated with secondary antibodies in 0.3% Triton X-100, 5% normal donkey serum in PBS for 2 hours at RT, and nuclei were stained with 300nM DAPI (Invitrogen). Slides were mounted using ProLong Antifade Diamond (Thermo Fisher). Images were taken using a Nikon AR confocal laser scanning microscope (Nikon).

### Live Cell Imaging

Live imaging was performed using the Nikon AR confocal laser scanning microscope equipped with temperature and CO2 control. For calcium imaging, Fluo-4 direct (Life Technologies) was prepared according to the manufacturer’s protocol. D83 organoids were dissociated using Accutase and plated on reduced growth factor Matrigel (Corning) coated 24 well plates. Cells were cultured for 1 week in maturation media with 48-hour media changes before dosing experiments were conducted. Organoids cells were incubated with Fluo-4 60 min prior to the start of imaging. Frames were taken every 8s for 10 minutes using a 10X objective. Following baseline measurement, plated cells were incubated in dopamine, either 1µM or 1mM, 1µM of D1 agonist SKF-81297 (Tocris Biosciences), or 1µM of Quinpirole (Tocris Biosciences) before a further round of imaging. For TTX (Sigma) measurements, organoids were incubated in 1µM TTX for 30 minutes following baseline measurements before another 10 minutes of imaging.

Data analysis was performed using FIJI. Regions of interest were manually selected using ROI manager, and mean fluorescence was calculated for each timeframe. Change in fluorescence was calculated as follows: ΔF/F =(F-F0)/F0, in which F0 was the mean fluorescence value recorded at t = 0, and a cell was considered electrically active if the fluorescence intensity change was greater than 0.2 ΔF/F. Cells were considered electrically active if they had at least one observable calcium spike during the 10-minute period. For figure 1 panel E, cells firing frequency (number of calcium transients per minute) and amplitudes (ΔF/F) were compared to their baseline firing frequency and amplitude before dopamine or ascorbic acid exposure. If there was a 20% increase in firing frequency or amplitude following compound exposure, cells were considered increased relative to baseline and if there was a 20% decrease cells were considered decreased relative to the baseline. For all measurements n=3 biological replicates of 3-5 dissociated and plated organoids were measured.

For cAMP imaging, the cADDis green down kit (Montana Biosciences) was used, and cells were prepared according to the manufacturer’s protocol. D83 organoids were dissociated using Accutase and plated on reduced growth factor Matrigel coated 24 well plates. Cells were cultured for 1 week in maturation media with 48-hour media changes before dosing experiments were conducted. Frames were taken every 8s for 20 minutes using a 10X objective. Dopamine was added at the 5-minute mark with the preceding 5 minutes considered the baseline measurement. For all ROS inhibitor and dopamine receptor antagonist measurements, plated cells were treated with either 100µM of acetylcysteine (Selleckchem) or 100µM of D1 antagonist SCH-23390 (Tocris Biosciences) and Sulpiride (Tocris Biosciences) for 30 minutes prior to imaging.

Data analysis was performed using FIJI. Regions of interest were manually selected using ROI manager, and mean fluorescence was calculated for each timeframe. Change in fluorescence was calculated as follows: ΔF/F =(F-F0)/F0, in which F0 was the mean fluorescence value recorded at t = 0. ΔF/F measurements were plotted over time revealing an s-curve when the change in fluorescence decreased. The amplitude change was considered the linear region of the s-curve, and the response time was calculated as the time after dopamine addition before the initial decrease in fluorescence intensity defined as the top of the linear section of the s-curve. For all measurements n=3 biological replicates of 3-5 dissociated and plated organoids were measured.

### Dosing Experiments

To perform acute and chronic dopamine dosing for live cell imaging. Plated D83 organoids were dosed with either 1µM or 1mM of dopamine in 100µM of ascorbic acid or a 100µM ascorbic acid control (Sigma) for 1 hour before the media was removed and replaced with fresh maturation media. For chronic samples, the plated cells were dosed with dopamine every day for 1 week before an acute dose was administered during live cell imaging. Acute samples and controls were dosed with ascorbic acid during the week leading up to imaging.

For bulk sequencing and immunohistochemistry dosing, D83 organoids were treated with either a dose of dopamine diluted in 100µM of ascorbic acid, cocaine (NIDA Drug supply program) diluted in 100µM of ascorbic acid, or 100µM ascorbic acid as a control for 1 hour before media was removed and replaced with fresh maturation media. Final concentrations of dopamine and cocaine were 1µM while the concentration of ascorbic acid was kept constant. For chronic samples, organoids were dosed daily for 1 week before a final 1-hour dose was administered. Acute samples and controls were dosed with ascorbic acid during the week prior before an acute dose of either dopamine or ascorbic acid was added. For all organoids treated with the ROS inhibitor, 100µM of acetylcysteine was added to the media prior to dosing with dopamine or cocaine. For the final dose following the 1-hour incubation, organoids were either immediately washed once with cold PBS and fixed for sectioning and immunostaining or washed twice with cold PBS before the samples were flash frozen in liquid nitrogen and shipped on dry ice for RNA extraction and sequencing.

### ROS Measurement

Oxidative stress experiments were modified from a previously described protocol^29^. Organoids were incubated with 100 μM 2′,7′-dichlorofluorescein diacetate (DCFH-DA) (SigmaAldrich) for 1 hour along with dopamine, cocaine, or ascorbic acid control. Organoids were washed twice in PBS before being dissolved in 1% Triton in PBS. Fluorescence measurements were taken at an excitation wavelength of 485nm and an emission wavelength of 530nm using a Tecan plate reader. Protein concentrations were obtained using the BCA assay (Thermo Fisher) according to the manufacturer’s instructions. ROS levels were calculated by dividing the fluorescence measurements by the total protein concentration. All ROS values were then normalized to the vehicle controls.

### RNA Extraction and Sequencing Analysis

Total RNA was extracted on site by the company performing sequencing. Sequencing and preparation of Illumina libraries was performed and then sequenced using Illumina Hi-seq 2 × 150 bp (Azenta, South Plainfield, NJ, USA or LC Sciences, Houston, TX, USA) for 20–30 million reads per sample. Raw sequencing data are publicly available on the Gene Expression Omnibus (GEO Accession Number:).

Raw FASTQ formatted sequence reads were imported into CLC Genomics Workbench (v.21.0.5 Qiagen, https://digitalinsights.qiagen.com/). Adaptor sequences and bases with low quality were trimmed and reads were mapped to the reference genome (GRCh38.102) using the RNA-seq analysis tool with the default parameters recommended for RNA-seq analysis. Principal component analysis and differential expression analysis were performed using ‘PCA for RNA-seq’ and ‘Differential Gene Expression for RNA-seq’ toolsets. Differential gene expression was performed by comparing organoids either acutely or chronically dosed with dopamine or cocaine against the ascorbic acid dosed controls. Genes were considered differentially expressed if they had a log2FC>|1| and a false discovery rate (FDR)<0.05.

For gene set enrichment analysis, all genes with a FDR<0.05 were uploaded into Ingenuity Pathway Analysis (IPA) software^82^ (Qiagen, https://digitalinsights.qiagen.com/IPA). Top regulated canonical pathways were predicted using -log10(p-value) while upstream regulators and comparisons were ranked based on z-score^82^. All plots and heatmaps were generated in Microsoft excel and figures were compiled in adobe illustrator. Overlaps between gene sets were performed using the Venn Diagram tool from the Bioinformatics & Evolutionary Genomics Department at Ghent University (https://bioinformatics.psb.ugent.be/webtools/Venn/).

### Statistical Analysis

Statistics were performed for live cell imaging and immunostaining quantification using the one-way ANOVA with Tukey’s post hoc test to test for significance. All analyses were performed using Microsoft Excel. Significance was defined as p < 0.05. Statistical analyses for RNA-seq were performed in CLC Genomics Workbench (v.21.0.5 Qiagen, https://digitalinsights.qiagen.com/). All sequencing data passed default quality filters for the CLC Genomic Workbench version 21.0.5 RNA-Seq pipeline analysis. Significance for Ingenuity Pathway (v. 22.0, QIAGEN Inc., https://digitalinsights.qiagen.com/IPA, accessed on 13 January 2022) and analyses were defined as q-value < 0.05 and z-score>|2|.

## Data Availability

The data can be accessed at: GSE234769.

## Acknowledgments

This work was funded by the NIH Avenir Award (DP1-DA044359) and the Ruth L. Kirschstein Research Award (F31 DA053128-01). Cocaine was provided by the National Institutes of Drug Abuse (NIDA) as part of their drug supply program. We thank Dilara Sen, Ryan Tam, and R. Chris Estridge for their thoughtful discussions and helpful insights.

## Author Contributions

T.R. and A.J.K. conceived the study. T.R. performed the wet lab experiments and research. T.R. performed bioinformatic analysis. T.R. and A.J.K. wrote the manuscript.

