## Supplemental Figures and Tables for "Reactive Oxygen Species Mediate Transcriptional Responses to Dopamine and Cocaine in Human Cerebral Organoids"

Thomas Rudibaugh, *et al.*

This file includes:
Figures S1-S4
Tables S1-S2

Not Included:
Supplemental Excel Files 1-4

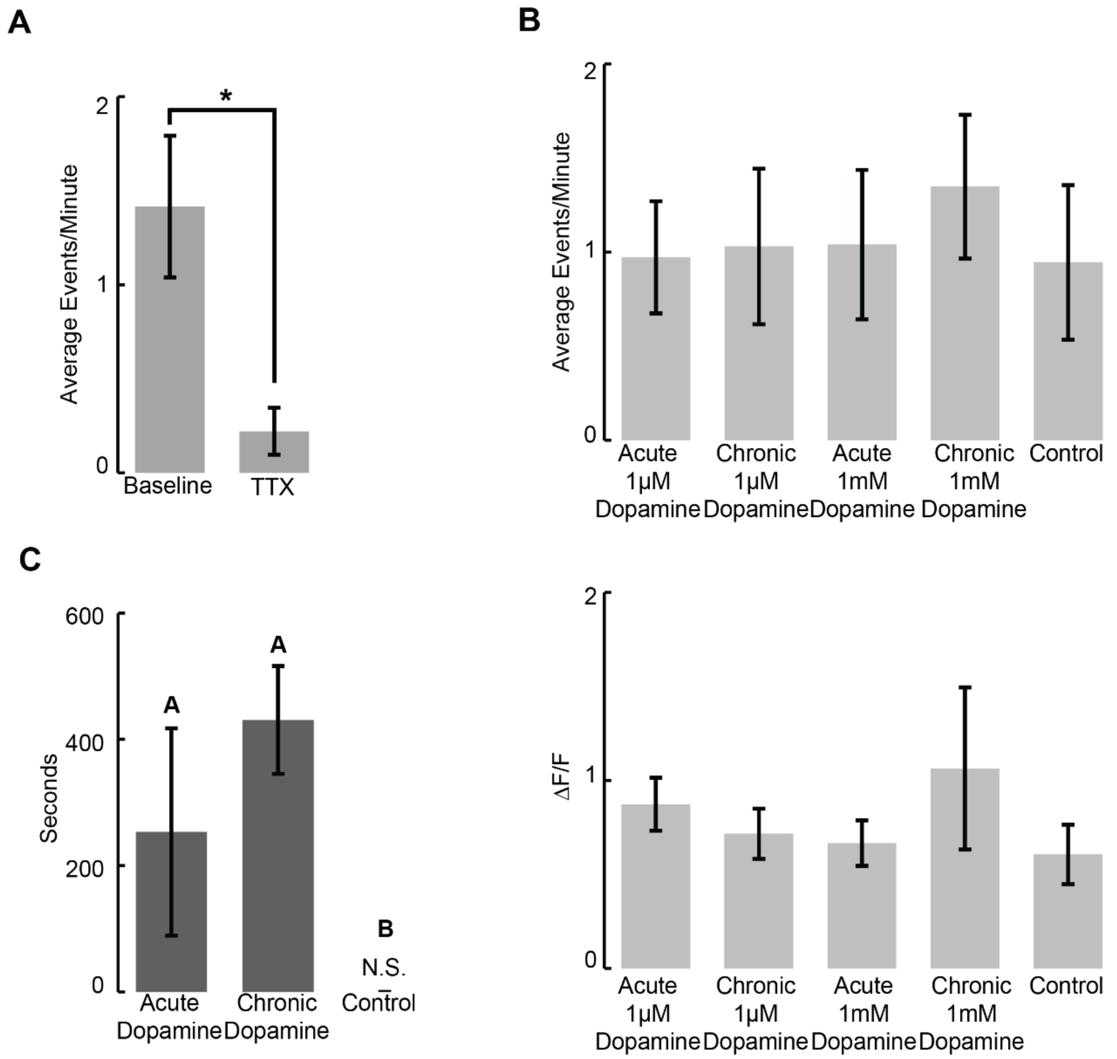


Figure S1, Related to Figure 1; ***Ventral forebrain organoids exhibit $Ca^{2+}$ and intracellular cAMP responses to dopamine***

**A)** $Ca^{2+}$ imaging of D90 organoids shows transients knocked down by TTX. Average transients/minute were measured for active cells (defined as any cell exhibiting at least one $Ca^{2+}$ transient during the measurement period). **B)** Frequency and amplitude of $Ca^{2+}$ transients following acute or chronic 1μM or 1mM dopamine exposure. Measurements were averaged across all active cells. **C)** Response time of intracellular cAMP increase following acute or chronic 1μM dopamine exposure. **A-C)** Error bars represent 95% confidence for n=3 replicates. **A)** * $p<0.05$, paired t-test. **C)** * $p<0.05$, one-way ANOVA with Tukey Kramer post hoc analysis.

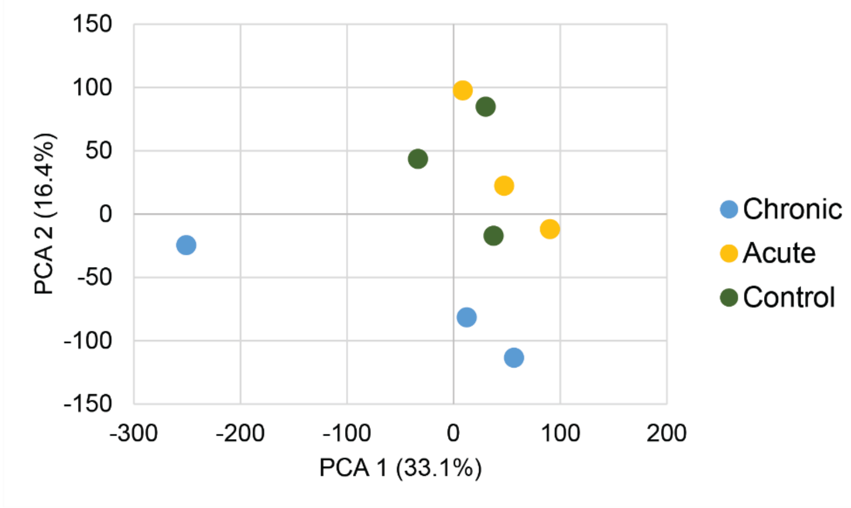


Figure S2, Related to Figure 2- ***Transcriptional analysis reveals immune-related response to acute and chronic dopamine***
Principal component analysis of the three biological replicates for the three different conditions: chronic dopamine, acute dopamine, and vehicle control.

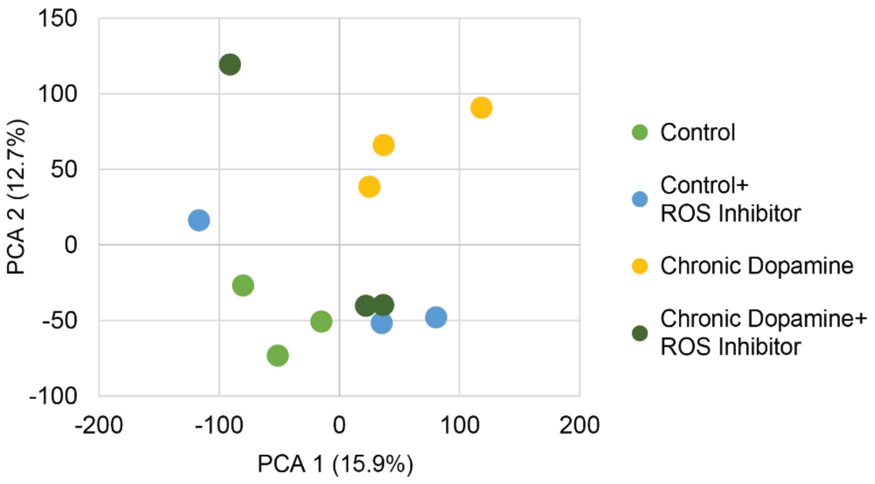


Figure S3; Related to Figure 3- ***Organoid response to dopamine driven by reactive oxygen species***
Principal component analysis showing the three replicates for the four conditions: control, control dosed with an ROS inhibitor, chronic dopamine dosed, and chronic dopamine dosed with an ROS inhibitor.

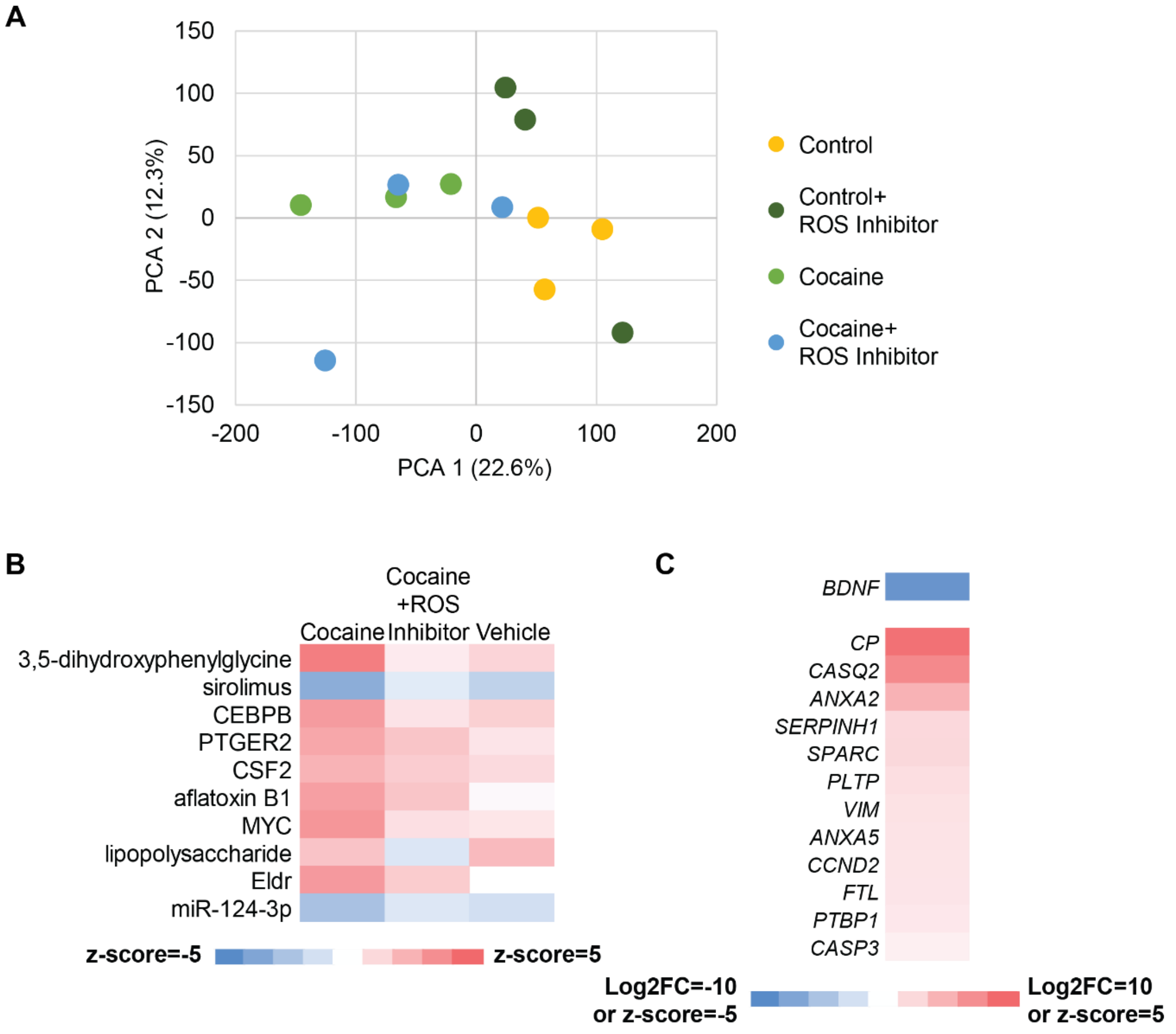


Figure S4; Related to Figure 4- ***ROS also mediates the transcriptomic response to cocaine***
**A)** Principal component analysis of the three biological replicates for the four dosing conditions: vehicle controls, vehicle controls with an ROS inhibitor, chronic cocaine, and chronic cocaine with an ROS inhibitor. **B)** Heat map of the top regulated upstream regulators by z-score identified using GSEA following dopamine or cocaine exposure with or without an ROS inhibitor. **C)** Heat map of BDNF predicted z-score and Log2FC of genes associated with BDNF downregulation based on Ingenuity Pathway Analysis (IPA) database.

**Table S1**
**Related to Figure 1, Primary Antibodies**

| Antigen | Species | Company | Catalog | RRID | Dilution |
|---|---|---|---|---|---|
| DARPP32 | Rabbit | Abcam | Ab40801 | AB_731843 | 1:50 |
| MAP2 | Mouse | Sigma-Aldrich | M1406 | AB_477171 | 1:250 |
| GABA | Rabbit | Sigma-Aldrich | A2052 | AB_477652 | 1:200 |
| CTIP2 | Rat | Abcam | Ab18465 | AB_2064130 | 1:250 |
| GSX2 | Rabbit | Millipore | Abn162 | AB_11203296 | 1:500 |
| SOX2 | Goat | R&D Systems | AF2018 | AB_355110 | 1:100 |

**Table S2**
**Related to Figure 1, Secondary Antibodies**

| Species | Flourophore | Company | Catalog | RRID | Dilution |
|---|---|---|---|---|---|
| Rabbit | 488 | Life Technologies | A21206 | AB_2535792 | 1:250 |
| Mouse | 546 | Thermofisher | A10036 | AB_2534012 | 1:250 |
| Rat | 647 | Jackson Immunoresearch | 712605150 | AB_2340693 | 1:125 |
| Goat | 647 | Jackson Immunoresearch | 705605003 | AB_2340436 | 1:125 |
